# Straintables: An application that extracts sequences from genome assemblies and generates dissimilarity matrices

**DOI:** 10.1101/2021.07.06.451382

**Authors:** Gabriel Nogueira Araujo, Richard W Francis, Cristina dos Santos Ferreira, Alba Lucínia Peixoto Rangel

## Abstract

**Background and Objectives:** The dissimilarity matrix (DM) is an important component of phylogenetic analysis, and many software packages exist to build and show DMs. However, as the common input for this type of software are sequences in FASTA file format, the process of extracting and aligning each set of sequences to produce a big number of matrices can be laborious. Additionally, existing software do not facilitate the comparison of clusters of similarity across several DMs built for the same group of individuals, using different genomic regions. To address our requirements of such a tool, we designed *Straintables* to extract specific genomic region sequences from a group of intraspecies genomic assemblies, using extracted sequences to build dissimilarity matrices.

**Methods:** A Python module with executable scripts was developed for a study on genetic diversity across strains of *Toxoplasma gondii*, being a general purpose system for DM calculation and visualization for preliminary phylogenetic studies. For automatic region sequence extraction from genomic assemblies we assembled a system that designs virtual primers using reference sequences located at genomic annotations, then matches those primers on genome files by using regex patterns. Extracted sequences are then aligned using Clustal Omega and compared to generate matrices.

**Results:** Using this software saves the user from manual preparation and alignment of the sequences, a process that can be laborious when a large number of assemblies or regions are involved. The automatic sequence extraction process can be checked against BLAST results using extracted sequence as queries, where correct results were observed for same-species pools for various organisms. The package also contains a matrix visualization tool focused on cluster visualization, capable of drawing matrices into image files with custom settings, and features methods of reordering matrices to facilitate the comparison of clustering patterns across two or more matrices.

**Conclusion:** *Straintables* may replace and extend the functionality of existing matrix-oriented phylogenetic software, featuring automatic region extraction from genomic assemblies and enhanced matrix visualization capabilities emphasizing cluster identification. This module is open source, available at GitHub (https://github.com/Gab0/straintables) under a MIT license and also as a PIPY package.

**Highlights:** Simple in-silico protocol for generation, visualization and comparison of dissimilarity matrices.

Accurate automatic sequence extraction from multiple genomic assemblies by using virtual primers built from reference sequences in an annotation file.

Draws matrices as images, with enhanced cluster visualization and customized options.

Supports reordering of matrix indices to better visualize clustering pattern conservation across multiple regions.

## Introduction

The graphical comparison of similarity for equivalent genomic regions across an intraspecies group of isolates is widely used in phylogenetic and parasitological studies (Sivakumar et al. 2019; Sridhar et al. 2019). Various computational tools have been created for this purpose, such as MatGAT (Campanella, Bitincka, and Smalley 2003) which is capable of creating identity matrices from multi-FASTA sequence files. Another way to represent phylogenetic data are dendrograms, generated by software like Clustal Omega (Sievers et al. 2011). DMs can be converted into dendrograms with algorithms such as UPGMA or neighbour joining, which can be done with the suite MEGA7 (Kumar, Stecher, and Tamura 2016). Articles featuring phylogenetic analysis often publish figures of dendrograms, not matrices. However, when the comparison of different genomic regions and their clustering patterns across different genomic regions within the same group of strains needs to be compared, dendrograms can be complicated to interprete, especially with larger individual counts.

When we had to compare clustering variation on several *Toxoplasma gondii* strains across several regions, it was not possible to find free software capable of drawing matrices with acceptable layout and looks for publication while also allowing for cluster visualization. We developed *Straintables* to address this issue. Among the novel matrix comparison methods it presents an engine that extracts sequences from genomic assemblies by using virtual primer pairs. This method automates the process of gathering the sequences that will be used to create DMs, and is useful for making a quick preliminary comparison of intraspecifc genomic regions of interest or an analysis involving a large number of regions, which would be laborious using conventional methods.

Recent advances in genomic sequencing technologies has resulted in a large number of genomic assemblies becoming available in public databases, were many species are represented by more than one assembly. While such volume of information is certainly useful for phylogenetic studies, it can be difficult to manage and prepare a large number of files to analyze. The automatic steps for sequence extraction from assemblies presented by *Straintables* can assist in analyses involving a large number of assemblies and regions, which would otherwise require substantial manual efforts or the design of ad hoc scripts to prepare input sequence files.

The source code of this application is available at GitHub (https://github.com/Gab0/straintables) under a MIT license. It takes as input a list of genomic regions of interest, a pool of intraspecies complete and/or partial genome assemblies, and an annotation file that corresponds to one of the assemblies. *Straintables* outputs various files, including matrix values files that can be viewed using a viewer which is included in the package. The Python language was chosen due to its fast development time and the vast library of modules available at pipy, from where we use Biopython (Cock et al. 2009), Scikit-bio, Flask, Matplotlib and others.

## Methods

### Protocol

The conversion of genetic input data into DM cell values involves a number of computational steps, where for each step this package provides a corresponding executable. In order to allow the execution of the full analysis with a single command, two pipelines were designed and are the recommended way to use the software. The first pipeline is the core *Straintables* method of extracting sequences from genomes and the other allows the user to provide the sequences that will build the DMs. Both pipelines will output equivalent files to the designated output directory, including multi-FASTA files with extracted sequences, alignment files, and DM values for each genomic region. With the output directory populated, the user can view the resulting matrices with the viewer application.

### Genomic Pipeline

This pipeline builds a DM for each region in the user provided list. In order to correctly extract region sequences from the pool of genomes, it reads both positional information for each region and the raw chromossome sequence contained in the annotation file to create template sequences that will be used to generate virtual primers to be be matched on the genomes. As an alternative to providing the assemblies and annotation file manually, a script to download genomes from the Assembly database at NCBI (Benson et al. 2012) is provided.

Once a pair of primers is randomly selected from the region template sequence, the program tries to match those primers in each genomic assembly. If both primers match in every assembly, the sequence between those primers, for all assemblies, is saved in a multi-FASTA file containing a track per assembly, with the extracted sequence. Otherwise, if any primer fails to match any assembly, the primer is discarded and a new primer is generated from the template region sequence. The maximum number of match failures is configurable. Figure 1a is a graphical representation of the steps of operation done by the genomic pipeline.

**Figure 1:**
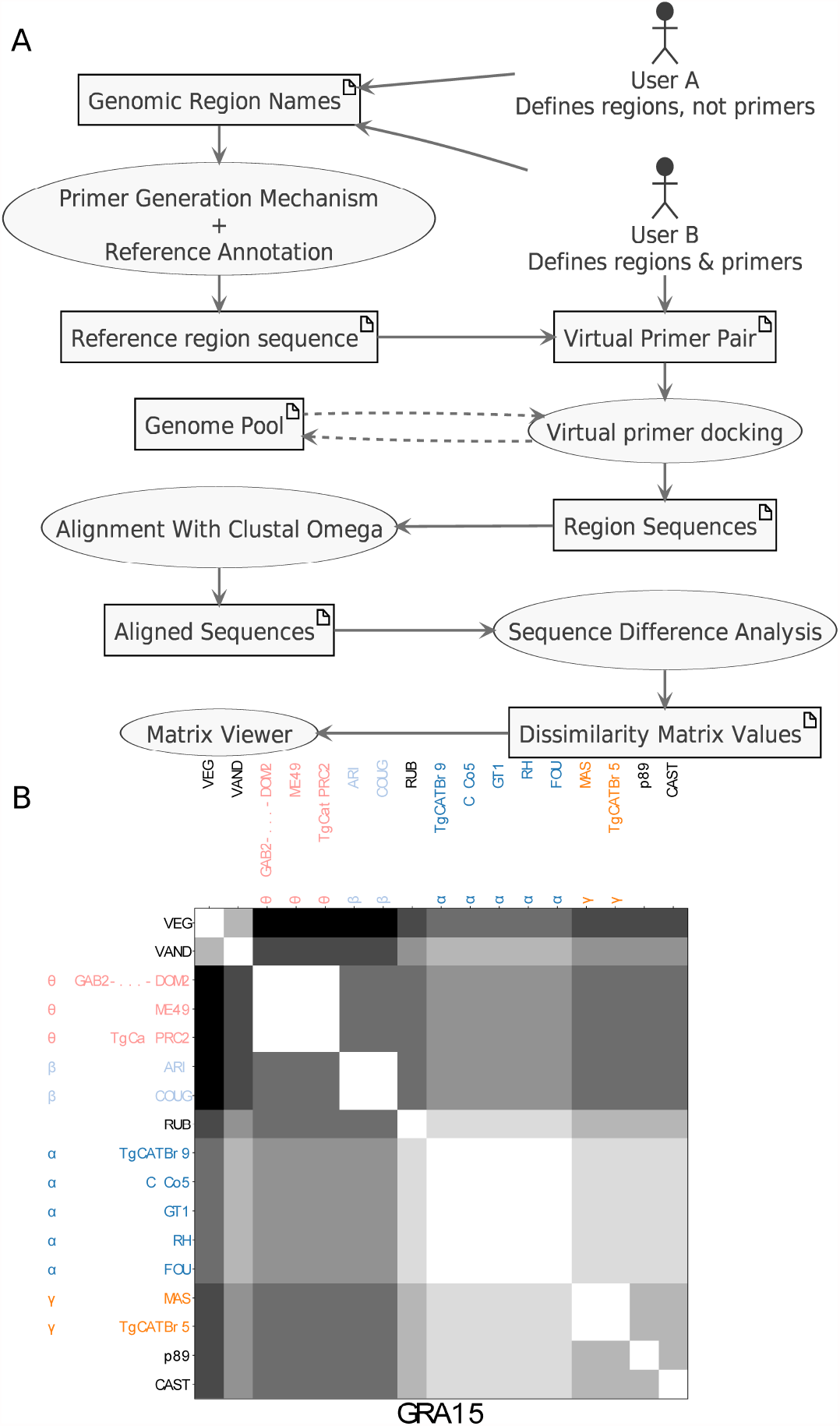
(a) Schematic of the genome pipleline workflow. In the image, each user represents a different choice of input. User A defines genomic region names but does not define primers, so the pipeline will launch the primer generation process trying to determine valid primer pairs for the desired regions. User B defines regions and primers, so provided primers will be matched on the genomes. Primer pairs are designed from region-specific sequences provided by the annotation file, and the genomic positions they match at each genome file are used as boundaries for region extraction. For each region, extracted sequences for all genomes undergo alignment and similarity comparison steps before the dissimilarity matrix for that region is drawn. (b) Dissimilarity Matrix built for the GRA15 region of *Toxoplasma gondii* genome. By using available genomes in NCBI database and the annotation for the ME49 strain, *Straintables*’ genomic pipeline found the forward primer 5’CTCAGGTAGTGATAGCCGGA3’ and the backward primer 5’CAGGGTCACGTACACAACCC3’. The software then extracted sequences between those primers for each assembly, aligned the sequences using Clustal Omega and built the DM using pairwise distances. 5

Next, DM cell values are calculated by counting SNPs shared between the pair of individual sequences at that specific column and row then dividing the number by the total number of SNPs found in the alignment. We can rely on such simple algorithm for distance calculation because the complex step of sequence alignment is done by Clustal Omega, which is a requirement for the genomic pipeline. Clustal Omega was chosen due to its reliable alignment results with default options and good performance.

The quality of genomic assemblies available in public databases vary, and certain assemblies may randomly split genomic regions into different multi-FASTA tracks, instead of the expected track for each chromosome. If a region is split into multiple tracks in the assembly it may not be possible to extract that region sequence, therefore the use of high quality genomic assemblies is recommended.

The sequences extracted from genomes by using this genomic pipeline can further be checked for correctness by doing BLAST (Altschul et al. 1990) searches using extracted sequences as queries. Expected BLAST results were observed for each tested intra-species genomic pools, from virus to protozoans.

### Matrix Visualization

The viewer executable is a Flask web server, designed to be accessed with any internet browser. It offers diverse methods of DM rendering, along with customizable figure sizes and resolution, and the possibility of including DM cell values into the matrices. Labels in DMs denotes the source organism of each sequence, and are colorized by similarity clustering across DM source sequences. The viewer also allows for the reordering of matrix columns and rows, enabling two or more matrices of different regions to be compared side by side, which greatly helps the visualization of clustering pattern variations across a group of regions. Matrices are rendered using the Matplotlib (Hunter 2007) Python module, and Figure 1b shows a DM produced by the *Straintables*’ viewer application. Each selected region will result in a single DM, where cells are shown as blocks colored in grayscale where white represents equality between two source sequences and black represents the maximum pairwise difference observed in the sequence pool. The fractional values used to render the DM can be optionally drawn at each cell. The number of simultaneous genome assemblies to compose a DM is limited by the output image size, as matrices built from larger genome pools will result in larger images if the matrix labels should remain legible. *Straintables* by default calculates a font size compatible to the genome pool size, while the user has the option to select a specific size for label text.

## Conclusion

While existing software can build dissimilarity matrices from genomic sequences, the *Straintables* package provides a unified and extensible workflow for phylogenetic studies centered in specific genomic regions. The software handles every step from the download of genomes to visualization of dissimilarity matrices, while also automating steps such as sequence preparation and alignment. This enables similarity distance comparisons of user defined genomic regions with minimum user effort. Automatic sequence extraction and alignment from genomic assemblies relieves the user from those laborious steps. Each step of data processing done by the application produces output files, which can be manually checked and analyzed with other software. Providing this software as a web service was considered, however as the sequence extraction process involves reasonable processing time, the idea was discarded due to practical reasons. Additional information for usage instructions and associated scripts can be found on the readme file inside the software repository.

## Acknowledgements

The authors declare no conflicts of interest. Thanks to Michał J. Gajda for proofreading the article. This research did not receive any specific grant from funding agencies in the public, commercial, or not-for-profit sectors.

## Summary

Straintables aims to provide a simple in-silico methodology for phylogenetic studies, with automatic steps that might be useful for biologists or medical doctors who do not have extensive bioinformatics training or do not have available time to manually prepare genomic sequences for analysis.

Instead of requiring the user to provide the sequences that will generate dissimilarity matrices, it offers a virtual primer matching system that extracts sequences from a group of partial or complete genome assemblies, requiring a single annotation file that matches any used assembly. The user defines a list of genomic regions and by comparing the resulting matrices and observed clusters of similarity, information about genetic variation across regions of multiple strains of an organism can be obtained.

The limit on the number of simultaneous genome assemblies to compose a DM is defined by how large the output image is allowed to be, with bigger genome pools producing larger images. In order to ensure clarity for label texts, the software calculates a default font size according to the genome pool size, while also enabling user-defined values.

This package is an open source Python module with executable scripts, offering a simple method for preliminary phylogenetic studies using dissimilarity matrices.

